# Amphibian segmentation clock models suggest mechanisms of slowed development across increasing genome size and nuclear volume

**DOI:** 10.1101/2023.07.16.549220

**Authors:** Alexandra Nicole Taylor, Rachel Lockridge Mueller, Ashok Prasad

## Abstract

Evolutionary increases in genome size, cell volume, and nuclear volume have been observed across the tree of life, with positive correlations documented between all three traits. Developmental tempo slows as genomes, nuclei, and cells increase in size, yet the driving mechanisms are poorly understood. To bridge this gap, we use a mathematical model of the somitogenesis clock to link slowed developmental tempo with changes in intra-cellular gene expression kinetics induced by increasing genome size and nuclear volume. We adapt a well-known somitogenesis clock model to two model amphibian species that vary ten-fold in genome size: *Xenopus laevis* (3.2 Gb) and *Ambystoma mexicanum* (32 Gb). Based on simulations and backed by analytical derivations, we identify parameter changes originating from increased genome and nuclear size that slow gene expression kinetics. We simulate biological scenarios for which these parameter changes mathematically recapitulate slowed gene expression in *A. mexicanum* relative to *X. laevis*, and we consider scenarios for which additional alterations in gene product stability and chromatin packing are necessary. Results suggest that slowed degradation rates as well as changes induced by increasing nuclear volume, which remain relatively unexplored, are significant drivers of slowed developmental tempo.

## Introduction

Across the tree of life, genome size and cell volumes span a remarkable range, and a positive correlation has been observed between increases in genome size, cell volume, and nuclear volume (Gregory 2001, Malerba and Marshall 2021, Sessions 2008). However, the mechanisms underlying these relationships and the implications associated with such increases remain areas of ongoing research. For example, evolutionary increases in genome, cell, and nuclear size have been found to slow developmental processes (Jockusch 1997, Sessions and Larson 1987, Wyngaard et al. 2005), but the driving mechanisms are poorly understood. Development emerges from the progression and interaction of a wide range of processes taking place at the single-cell level, where increasing genome, cell, and nuclear size impact transcription dynamics, intra-cellular distances, surface area to volume ratios, and other fundamental characteristics (Cadart et al. 2023, Sessions and Wake 2021). We therefore consider how alterations in single-cell processes might translate to slowed developmental tempo as increasing genome, cell, and nuclear sizes change cell structure and functionality.

Vertebrates comprise a large portion of the overall range in genome and cell size across the tree of life, and despite variation in genome size, cell volume, and developmental tempo, many developmental processes remain conserved. Somitogenesis is one such process that is relatively well understood; it is the process through which bilateral pairs of somites, blocks of presomitic mesoderm (PSM) tissue, are patterned along the head-to-tail axis in vertebrate embryos. Somitogenesis is typically described as operating through a clock and wavefront mechanism in which cell-autonomous oscillations of a somitogenesis gene (i.e. the “segmentation clock”) interact with Notch, Wnt, FGF, and retinoic acid pathways across the PSM tissue to coordinate proper timing of segmentation of groups of neighboring cells into bilateral pairs of somites (Aulehla and Pourquié 2010, Cooke and Zeeman 1976, Gibb et al. 2010, Klepstad and Marcon 2023). The segmentation clock operates via oscillatory gene expression at the single-cell level, while the wavefront takes place at the intercellular level across the PSM. Vertebrates exhibit species-specific segmentation clocks directly related to oscillatory expression of an autoregulated somitogenesis gene at the single cell level (Diaz-Cuadros et al. 2023, Lázaro et al. 2023, Matsuda et al. 2020). One period of oscillation, the time required for one cycle of expression, at the single cell level determines the time needed for a group of neighboring cells to segment off from the larger block of unsegmented PSM tissue, creating a bilateral pair of somites. As a conserved phenomenon that drives developmental tempo while operating at the single-cell level, the segmentation clock is an appropriate lens through which to examine single-cell processes as an underlying link between increasing genome and cell size and slowed development. While species-specific segmentation rates have been linked to biochemical differences at the intracellular level (Diaz-Cuadros et al. 2023, Lázaro et al. 2023, Matsuda et al. 2020, Rayon et al. 2020), the role of genome and cell size as potential mediators of species-specific gene expression kinetics and therefore developmental tempo have not been explicitly examined.

The segmentation clock is well modeled by the following system of delayed differential equations, first proposed by Lewis [2003], that describe coupled oscillatory expression of mRNA and protein,

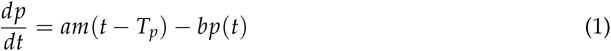

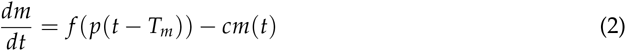

where: *p* is protein expression (i.e. the number of protein molecules in a cell); *m* is mRNA expression (i.e. the number of mRNA molecules in a cell); *a* is a rate constant for protein synthesis (protein/mRNA/min); *b* is the degradation rate for protein (protein/minute) and is given by the following expression: *b* = *ln*(2)/*h_p_*, where *h_p_* is the half-life (minutes) of the protein molecule; *c* is the degradation rate for mRNA (mRNA/minute) given by the following expression: *c* = *ln*(2)/*h_m_*, where *h_m_* is the half-life of the mRNA molecule; and *T_p_* and *T_m_* are the delays associated with protein and mRNA production, respectively. Parameter notation and descriptions are provided again in Table 1.

Equation (1) models the change in protein expression over time, 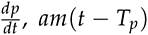 gives the rate at which proteins emerge in the cytoplasm, and *bp*(*t*) gives the rate at which they degrade. Similarly, equation (2) models the change in mRNA expression over time, 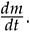. *cm*(*t*) describes the rate at which mRNA molecules degrade while *f* (*p*), given below, is a Hill function that describes the rate at which mRNA molecules emerge from the nucleus into the cytoplasm,

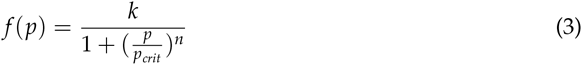

where *k* is a rate constant for mRNA synthesis (mRNA/min/cell) in the absense of repression, *p_crit_* is the number of protein molecules in the nucleus needed to yield an assumed critical concentration of 10*^−^*^9^ *M* within the nucleus (from Lewis [2003]) associated with transcriptional repression, and *n* is the so called “Hill coefficient”. The rate of mRNA emergence is dependent upon and inversely related to protein quantity, indicating the presence of an auto-repressive mechanism (i.e. a negative feedback loop). Like Lewis [2003], we assume that these repressive proteins act as dimers and let *n* = 2.

**Table 1:** Model parameters.

| Notation | Description |
| --- | --- |
| $p_{crit}$ | Critical protein threshold, number of protein molecules needed to achieve $10^{-9}$ concentration in nucleus |
| $T_p$ | Delay associated with protein production (min) |
| $T_m$ | Delay associated with mRNA production (min), sum of: |
| $T_{tx}$ | Transcription delay (min) |
| $T_{in}$ | Intron splicing delay (min) |
| $T_{exp}$ | mRNA export delay (min) |
| $a$ | Rate of protein synthesis (protein/mRNA/min) |
| $k$ | Rate of mRNA synthesis, no repression (mRNA/min/cell) |
| $b$ | Rate of protein degradation (protein/min), equal to $\ln(2)/h_m$ |
| $h_m$ | Protein half-life |
| $c$ | Rate of mRNA degradation (mRNA/min), equal to $\ln(2)/h_p$ |
| $h_p$ | mRNA half-life |

As touched on above, the parameters *T_p_* and *T_m_* account for delays associated with protein and mRNA production, respectively. Delay parameters reflect the reality that biological processes are often a non-instantaneous affair. Mathematically, they distinguish the DDE system above from an ordinary differential equation system and are necessary to generate the sustained oscillations that yield species-specific periods of gene expression (Lewis 2003), corresponding to species-specific rates of somite segmentation.

Here, we adapt Lewis’ model to assess the impact of increasing genome and cell size on the segmentation clock periods associated with two amphibian species that exhibit a 10-fold difference in genome size: *Xenopus laevis* and *Ambystoma mexicanum*, the model frog and salamander, respectively. We break delay parameters down into specific transcription, post-transcription, and translation processes, and we consider the potential impact of increasing genome and/or cell size on each individual component. We also adjust critical protein threshold values to reflect species-specific nuclear volume estimates. Finally, we consider additional potential roles for mRNA and protein stability, which are not directly related to genome or cell size, in the mediation of developmental tempo. We simulate the Lewis model under all of these different scenarios to test whether we can reproduce the observed periodicity of the somitogenesis clock. We buttress our simulations with an analytical derivation of the minimal conditions for oscillations in the Lewis model. With our approach, we are able to establish direct links between increases in genome size and nuclear volume and their specific impacts on gene expression (i.e. increases in network delays and threshold value) that yield slower developmental tempo, and we are also able to assess potential indirect roles for gene product stability in the mediation of developmental tempo.

## Methods

### Taxon selection

We choose to adapt the Lewis model to *X. laevis* and *A. mexicanum*. Their recorded genome sizes are *∼*3.1 Gb (*X. laevis*) and *∼*32 Gb (*A. mexicanum*) (Hellsten et al. 2010, Smith et al. 2019), and, although volumes vary across type, *A. mexicanum* cell volumes are typically larger than their *X. laevis* counterparts. *A. mexicanum* nerve cells, for example, are *∼*2-times larger than *X. laevis* nerve cells (Roth and Walkowiak 2015), and their red blood cells (RBCs) are *∼*10-times than in *X. laevis* (Gregory 2023, using cell volume measurements for *Ambystmoma tigrinum* whose average reported genome size is also *∼*32 Gb. Note also that amphibian RBCs are nucleated). Meanwhile, there is about a 3-fold difference in the rate of somite segmentation. In *X. laevis*, bilateral pairs of somites are segmented off from the PSM every 50 minutes (extrapolated by Curran et al. [2014] from Faber and Nieuwkoop [1994] and Hamilton [1969]); in *A. mexicanum*, somite segmentation occurs every *∼*155 minutes (Armstrong and Graveson 1988).

### Generating species-specific parameter values

Our first goal was to test if parameter changes directly related to increasing genome, cell, and nucleus size are sufficient to recapitulate the slowed rate of somite segmentation in *A. mexicanum* relative to *X. laevis*. To this end, we start by generating species-specific delay time and critical threshold parameters that capture genome size and nuclear volume, while holding all other parameters (rates of production and degradation) at constant values or ranges between species.

We derive protein and mRNA-production delays, *T_p_* and *T_m_*, by applying estimation methods from Lewis to somitogenesis gene candidates for *X. laevis* and *A. mexicanum*. Vertebrate clock genes are members of the Hairy and enhancer of Split (*Hes/Her*) family of basic helix-loop-helix (bHLH) genes (Kageyama et al. 2007). *Hes/Her* gene family size varies across vertebrates, and the individual cycling members of the clock network also vary (Eckalbar et al. 2012, Krol et al. 2011). In zebrafish and mice, *hes7* has been shown to be the central component of the oscillator (Bessho et al. 2001, Hirata et al. 2004, Holley et al. 2002, Oates and Ho 2002), and its oscillatory pattern has also been shown in the PSM in the lizard *Anolis carolinensis* (Eckalbar et al. 2012). We therefore infer that *hes7* is the ancestral clock for vertebrates and that it retains clock function in *A. mexicanum*. In *Xenopus laevis*, in contrast, the *Hes/Her* gene family is greatly expanded to 37 copies, following both ancient allotetraploidization and tandem duplication (Kuretani et al. 2021, Watanabe et al. 2017). The *hes7* orthologs in *X. laevis* are not cyclically expressed in the PSM and, therefore, cannot act as the clock (Davis et al. 2001, Jen et al. 1999, Shinga et al. 2001). In contrast, in *Xenopus laevis*, three *Hes/Her* family genes are known to oscillate in the PSM: *hes5.3*, *hes5.5*, and *hes5.7* (Blewitt 2009, Li et al. 2003). Of these, the strongest candidate is *hes5.7L* based on experimental data showing that *de novo* protein synthesis is required to repress *hes5.7L* transcription during somite formation, and that *hes5.7L* RNA instability is part of the mechanism underlying its cyclic expression (Davis et al. 2001, Li et al. 2003). *Hes5.7* is absent from the closely related *X. tropicalis* and is thus inferred to be specific to *X. laevis* (Kuretani et al. [2021], Watanabe et al. [2017]), suggesting a case of developmental system drift (Haag and True 2021). In *X. laevis*, *hes5.7L* has a primary sequence length of 1,604 nt; it is made up of 3 exons and 2 introns (lengths: 166 and 113 nt), and its coding sequence is 465 nt (Sayers et al. 2022). In *A. mexicanum*, *hes7* has a primary sequence length of 8272 nt; it is made up of 4 exons and 3 introns (lengths 3017, 1260, and 2030 nt), and its coding sequence is 783 nt (Smith et al. 2019 visualized on http://genome.ucsc.edu/). This information is used to derive species-specific parameters, described below:

#### Critical protein threshold, p_crit_

We calculate the number of protein molecules needed to achieve a critical concentration of 10*^−^*^9^ *M* in the nucleus, based on species-specific nuclear volumes. We assume a spherical nucleus (*V* = ^4^ *πr*^3^) and estimate species-specific radii of 4 and 5.5 *µm* using Fiji analyses (Schindelin et al. 2012) of stained PSM nuclei in *X. laevis* (Hidalgo et al. 2009) and *A. mexicanum* (Banfi et al. 2012), respectively.

#### Protein production delay, T_p_

We assume that the delay associated with protein production is equal to translation delay. We estimate species-specific translation delays by applying a translation rate of 6 nucleotides per second (Lewis 2003) to the reported coding sequence lengths.

#### mRNA production delay, T_m_

We consider mRNA-production delay to be a cumulative sum of transcription *T_tx_*, intron-splicing *T_in_*, and mRNA export *T_exp_* delays. We estimate species-specific *T_tx_* values by applying a transcription rate of 20 nucleotides per second (Lewis 2003) to the reported primary sequence lengths. Hoyle and Ish-Horowicz [2013] find that in vivo intron-splicing delay constitutes a relatively constant proportion of *∼*8.3 % of the overall segmentation clock period in mice, chick, and zebrafish embryos. We apply this proportion to the reported clock periods in *X. laevis* (*∼*50 min) and *A. mexicanum* (*∼*155 min) to get species-specific *T_in_* values. We estimated species-specific mRNA export delays, *T_exp_*, using simulations of particle diffusion within a sphere, described in detail below.

#### Simulation of mRNA export time parameter, T_exp_

Before they are released into the cytoplasm, newly transcribed mRNA molecules must journey from their chromatin address to the nuclear periphery where they locate an exit pore. Journeying to the periphery constitutes a relatively large part of this process, on the order of minutes, whereas locating and exporting through a pore once there is relatively rapid, typically on the order of fractions of seconds (Ben-Yishay and Shav-Tal 2019, Mor and Shav-Tal 2010, Mor et al. 2010). mRNA movement through the nucleoplasm takes place via passive diffusion. Both normal and obstructed (sub-) diffusion have been observed (Ben-Ari et al. 2010, Ishihama and Funatsu 2009, Mor and Shav-Tal 2010, Mor et al. 2010, Oeffinger and Zenklusen 2012, Shav-Tal et al. 2004). Normal and obstructed diffusion of RNA can be described by simple and fractional Brownian Motion, respectively (Jeon et al. 2014, Lampo et al. 2017, Mor and Shav-Tal 2010, Oeffinger and Zenklusen 2012), with obstructed diffusion in the nucleus typically attributed to constrained pathways arising from chromatin organization (Ben-Ari et al. 2010, Ishihama and Funatsu 2009, Mor and Shav-Tal 2010, Oeffinger and Zenklusen 2012, Sheinberger and Shav-Tal 2013). Given that both normal and obstructed diffusion of transcripts in the nucleus have been observed, we run normal and fractional Brownian Motion simulations and compare results.

To generate species-specific estimates for nuclear export, we simulate the (3D) random walk of a diffusing particle within spheres of radii between 3 and 6 *µ*m. This range of radii captures measurements for *X. laevis* and *A. mexicanum* PSM cell nuclei, 4 and 5.5 *µ*m, respectively. We simulate mRNA transcript trajectories and record the number of steps needed for our simulated mRNA molecule to first cross the nuclear periphery from the nuclear center. 10,000 trajectories are simulated for each sphere in the specified range (radius of 3 to 6 with intervals of 0.5).

The transcript trajectory for normal diffusion is simulated by generating x, y, and z position vectors as cumulative sums of increments (i.e. step sizes) chosen from a normal random distribution. The transcript trajectory for obstructed diffusion is simulated similarly, with the x, y, and z position vectors generated directly by a fractional Brownian Motion function in MATLAB, wfbm, with a Hurst parameter of 0.25. For both simulations, the number of steps required by a particle trajectory to first exit the domain is retrieved and averaged. We assume that each step takes one second, and we scale the particle trajectory such that the estimated mean first exit times agree reasonably well with mRNA exit times observed for the somitogenesis gene in *Danio rerio*, or zebrafish. It has been reported that nuclear export of *her1*(*hes7*), the somitogenesis clock gene in zebrafish, takes 3.36 minutes (Hoyle and Ish-Horowicz 2013). We used Fiji (Schindelin et al. 2012) and images of stained nuclei across the PSM in zebrafish from Keskin et al. [2018] to estimate a corresponding nuclear radius of 3*µ*m. Then, we scaled our trajectory time scales in both simulations such that it takes, on average, about 202 seconds, i.e., 3.36 min (corresponding to 202 steps of the simulation) to first exit a sphere of radius 3*µ*m.

We assume the nuclear center as the initial mRNA position based on known chromosomal territories associated with the somitogenesis gene in humans and mice coupled with patterns of synteny observed across vertebrates. That is, in both mice and humans, the somitogenesis gene, *hes7*, is found on gene-rich chromosomes 11 and 17 (which are homologs), respectively; both chromosomes localize in their respective nuclear centers Boyle et al. 2001, Kile et al. 2003, Mayer et al. 2005, Zody et al. 2006. Broad patterns of synteny conservation and topologically associated domain conservation have been observed across vertebrates (Schloissnig et al. 2021, Smith et al. 2019), suggesting that this pattern extends beyond humans and mice.

#### Parameters held constant

For the first set of analyses, we hold the rates of mRNA production (in the absence of inhibition), protein production, and mRNA degradation constant, at *k* = 33 mRNA/min, *a* = 4.5 protein/mRNA/min, and *c* = *ln*(2)/3 mRNA/min corresponding to a halflife *h_m_* = 3 minutes (Lewis 2003); and we consider a set range of protein stability/degradation *ln*(2)/23 *< b < ln*(2)/3 protein/min, corresponding to a half-life of 3 *< h_p_ <* 23 minutes. This range is chosen based on typical reported and estimated protein stability of the somitogenesis gene across model vertebrates, namely zebrafish and mice; we test the model across a range of protein half-lives due to its important role in mediating the period of gene expression (Hirata et al. 2004, Lázaro et al. 2023, Lewis 2003, Matsuda et al. 2020, Takashima et al. 2011) (see also Table 3: Sensitivity Analysis).

### Extrapolation of periodicity to compare with known somite segmentation rates

For our first set of analyses, we plug the parameter values described above back into the Lewis model, and we use the DDE solver ddesd in Matlab to generate solutions across identical ranges of protein stability *h_p_* and species- and diffusion-specific ranges of total delay time, *T_m_* + *T_p_*. We assess the period of gene expression that emerges for each set of solutions, and we compare it to the known somite segmentation rate of the corresponding species. In doing so, we aim to first verify that our parameter selection does in fact yield the correct period of oscillation for *X. laevis*, and to then determine if parameter changes directly driven by genome and cell size differences are sufficient to capture slowed developmental rate in *A. mexicanum* relative to *X. laevis*.

To assess periodicity, we create vectors to store the local extrema (i.e. local minimum and maximum values) and corresponding time stamps for each solution. Gene expression tends to spike in the first 4 to 5 cycles of oscillation before settling into a long-term pattern, so we remove the first 5 cycles of oscillation from our data to avoid skewing. The period of oscillatory gene expression is calculated by taking the average difference between successive time stamps associated with local minima (using local maxima would yield the same results). We do this across a time span of 0 to 3,100 minutes. The time span, 3,100 minutes, is chosen based on the fact that (at least) 20 somites are observed in *A. mexicanum* embryos, each requiring *∼*155 minutes to form (Armstrong and Graveson 1988). This time span also works well for *X. laevis*, with approximately 50 somites (Dali et al. 2002), each taking *∼*50 minutes to form (Li et al. 2003). For both species, oscillations must remain robust throughout this time span. We define robust oscillations as having an amplitude (the height of an oscillation, or difference between local minimum and maximum) of no fewer than 10 molecules throughout the time span. If this requirement is not met, the oscillations are considered damped and we define the periodicity as Inf (infinite). We choose 10 molecules as a conservative finite cut-off based on observed average RNA transcript amplitudes of zebrafish segmentation clock genes *her1* and *her7* which are *∼*41 and *∼*49 molecules, respectively (Keskin et al. 2018).

The assessment of periodicity described above is done for both mRNA and protein counts, and the difference in periodicity is always within 0.2 minutes (see Supplemental Material 1). In other words, results are relatively similar for both sets of oscillations, so we choose mRNA periodicity to represent overall system behavior.

### Calculating average amplitude of expression

We also calculate the average amplitude of mRNA expression across the parameter combinations in each model. To do this, we use the extrema vectors described above to create a vector that stores the difference between local minima and maxima corresponding to every complete cycle of oscillation, excluding data from the first 5 oscillation cycles that were cut out. The resulting vector gives the amplitude associated with each complete oscillation cycle, and we take the average of all vector entries to get an average amplitude associated with a particular parameter combination. For all combinations with periodicity defined as Inf (infinite), we set amplitude equal to 0, to keep results consistent with each other.

### Sensitivity analysis

To assess the impact of individual changes on the period of gene expression, we increase and decrease each individual parameter (assessing total delay as a parameter as opposed to taking *T_p_* and *T_m_* individually) by 50% while holding all other parameters constant. We first extrapolate the resulting period and amplitude of gene expression for an original set of parameters (corresponding to an *A. mexicanum* Brownian Motion model with protein half-life arbitrarily set at *h_p_* = 15 minutes) and then for each parameter change using the methods described above.

### Testing whether scaling mRNA stability with export time yields the rate of somite segmentation in A. mexicanum

Our mRNA export simulations suggest that transcripts take much longer to leave larger nuclei. We therefore assess the impact of increasing mRNA stability with estimated nuclear export time on the *A. mexicanum* segmentation clock period. To this end, we re-considered both *A. mexicanum* models, normal and fractional Brownian Motion corresponding to normal and obstructed diffusion, under 3 different levels of mRNA stability: half-life equal to mRNA export time, half-life equal to 50% of mRNA export time, and half-life equal to 25% of mRNA export time. While holding all other *A. mexicanum* species- and diffusion-specific parameters at their initial values (Table 2), we re-generate solutions and assess periodicity for the Lewis model given *h_m_* = *T_exp_*, 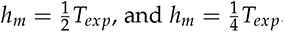.

**Table 2:**
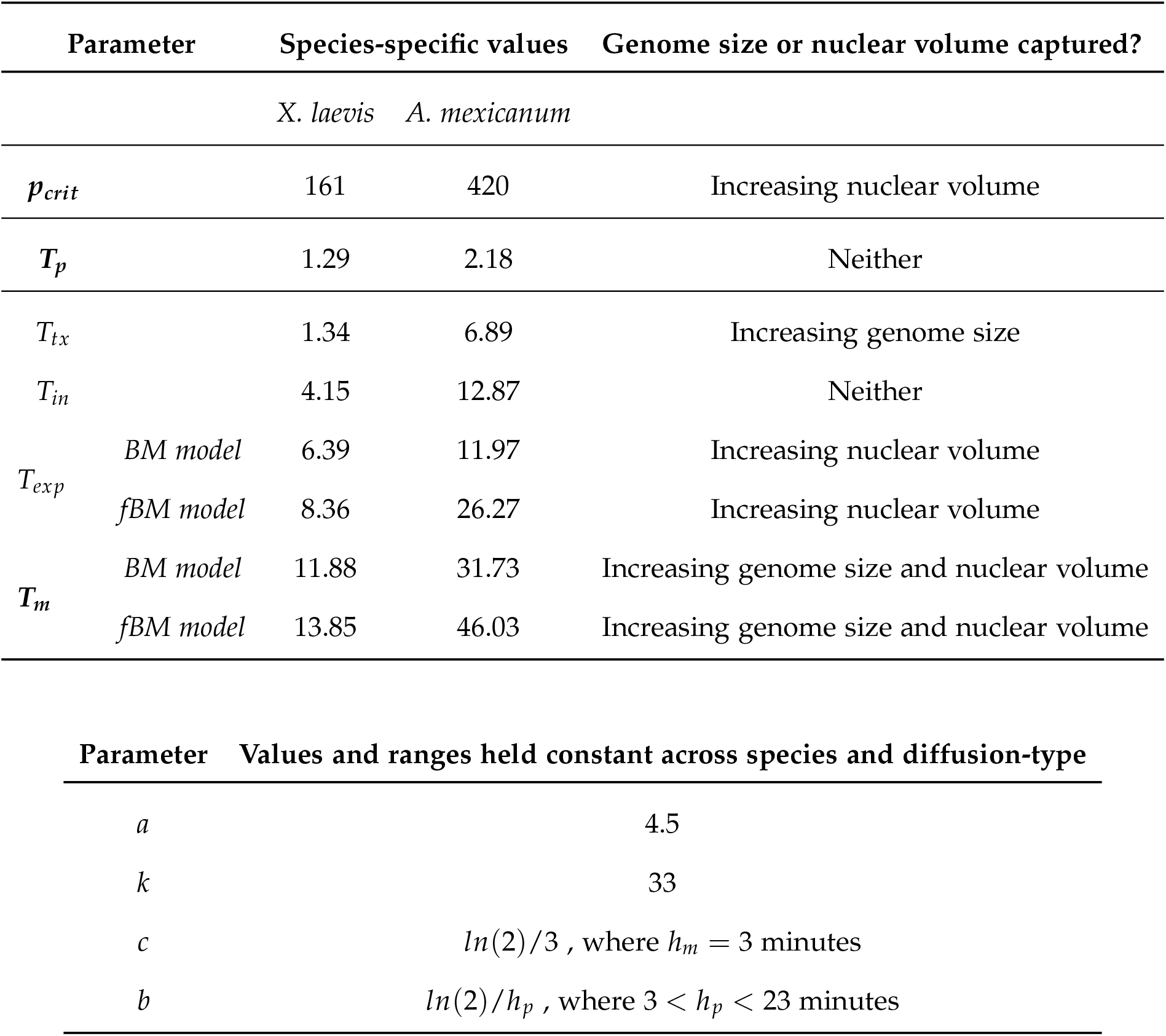
Initial parameter values and what is captured by changes in species-species values.

### Generalizing mRNA export simulations

We generalize our mRNA export simulations by running the simulations described above across a range that encompasses what has been observed across the tree of life, that is nuclei of radius between *∼*0.5 and 13 *µ*m, based on the minimum and maximum nuclear volumes reported in the dataset used by Malerba and Marshall [2021] (https://doi.org/10.5061/dryad.vq83bk3ss) while assuming a spherical volume 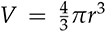. We also run simulations for which the initial position is selected from a uniform distribution as opposed to always being set at the origin (assumed to be at the nuclear center). The uniform distribution we draw our initial x, y, and z positions from encompasses a domain that is 3/4 of each radius. In doing so, we allow for initial positions to be drawn from positions throughout our theoretical nuclei, excluding the periphery. We choose to exclude the nuclear periphery because this is where heterochromatin is spatially concentrated and reduced transcriptional activity has been observed (Bizhanova and Kaufman 2021).

## Results

### Period of gene expression is most sensitive to changes in delay and stability parameters

Conditions for the emergence of oscillations for simplified versions of the Lewis model, assuming a single degradation rate and a single delay, have been obtained analytically Verdugo and Rand 2008, but it was not clear whether the results would generalize. We therefore derived conditions for the emergence of oscillations for the full Lewis Model (see Supplemental Material 2). We found that oscillating solutions require (i) the geometric mean of the degradation rates to be less than an upper bound, √K, and (ii) total delay *T_m_* + *T_p_* to be equal to or greater than a critical value, *T_crit_*. Both √*K* and *T_crit_* depend upon the kinetic constants of the model. In particular, *T_crit_* is positively correlated with *h_p_*, the protein half-life. Thus sustained oscillations with longer lived proteins require longer delay times.

Since analytical solutions of the Lewis model for arbitrary parameters do not exist, we simulated the model for different parameter values and calculated the period of oscillation. In agreement with results in Lewis [2003], we find that the period of gene expression (we look at mRNA specifically) is most sensitive to changes in total delay, *T_m_* + *T_p_*, and stability, *h_m_* and *h_p_*, parameters. Additionally, we find that when protein is too stable and total delay time is too low, robust oscillations do not emerge. In other words, total delay must be relatively large compared to protein half-life for oscillations to emerge, in agreement with our analytical results and previous mathematical and empirical results Hirata et al. 2004, Lewis 2003, Takashima et al. 2011.

### Parameter changes directly linked to increasing genome and cell size can mathematically recapitulate slowed developmental tempo

Using simulated export times, we generate species- and diffusion-specific general delay times associated with mRNA production, a sum of transcription, intron splicing, and nuclear export delays. All other parameters are held constant either across species (*a*, *k*, *b*, *c*) or at species-specific values across diffusion models (*p_crit_*, *T_p_*). All of these values are shown in Table 2.

The resulting periods of oscillation for each model are shown in figure 1 (with non-oscillatory combinations/regions, assigned period Inf, shown in dark purple). We test across a range of parameter combinations: on the x-axis, we have values of total delay, *T_m_* + *T_p_*, that fall within *±*5 minutes of our species- and diffusion-specific values given in Table 2; on the y-axis, we have a range of protein stability corresponding to 3 *< h_p_ <* 23 minutes. The observed rate of somite segmentation in *X. laevis* (*∼*50 minutes) is captured by a subset of total delay and protein stability combinations under both normal and sub-diffusive (obstructed diffusion) conditions; this subset is outlined in the plot by a dashed-line in figures 1A and 1B. In confirming that we can achieve the correct period of oscillation for *X. laevis*, the results in figures 1A and 1B provide support for our methods of parameter estimation; these results also act as a plausible baseline against which the *A. mexicanum* models can be compared. Meanwhile, the observed rate of somite segmentation in *A. mexicanum* (*∼*155 min) is only captured under sub-diffusive conditions. Genome and nucleus size-driven increases in delay time and concentration threshold are sufficient to fully recapitulate slowed development in *A. mexicanum* when nucleoplasmic movement of transcripts is assumed to be sub-diffusive (figure 1D), but are insufficient when normal diffusion is assumed (figure 1C). When normal diffusion is assumed, genome and cell size-driven parameter changes can slow the period of oscillatory gene expression down to *∼*125 minutes, 30 minutes faster than the known rate of somite segmentation. The additional delay introduced by assuming sub-diffusion, about 14 minutes longer, is needed to produce a set of total delay and protein stability combinations that yield a *∼*155 min period of oscillation.

**Figure 1:**
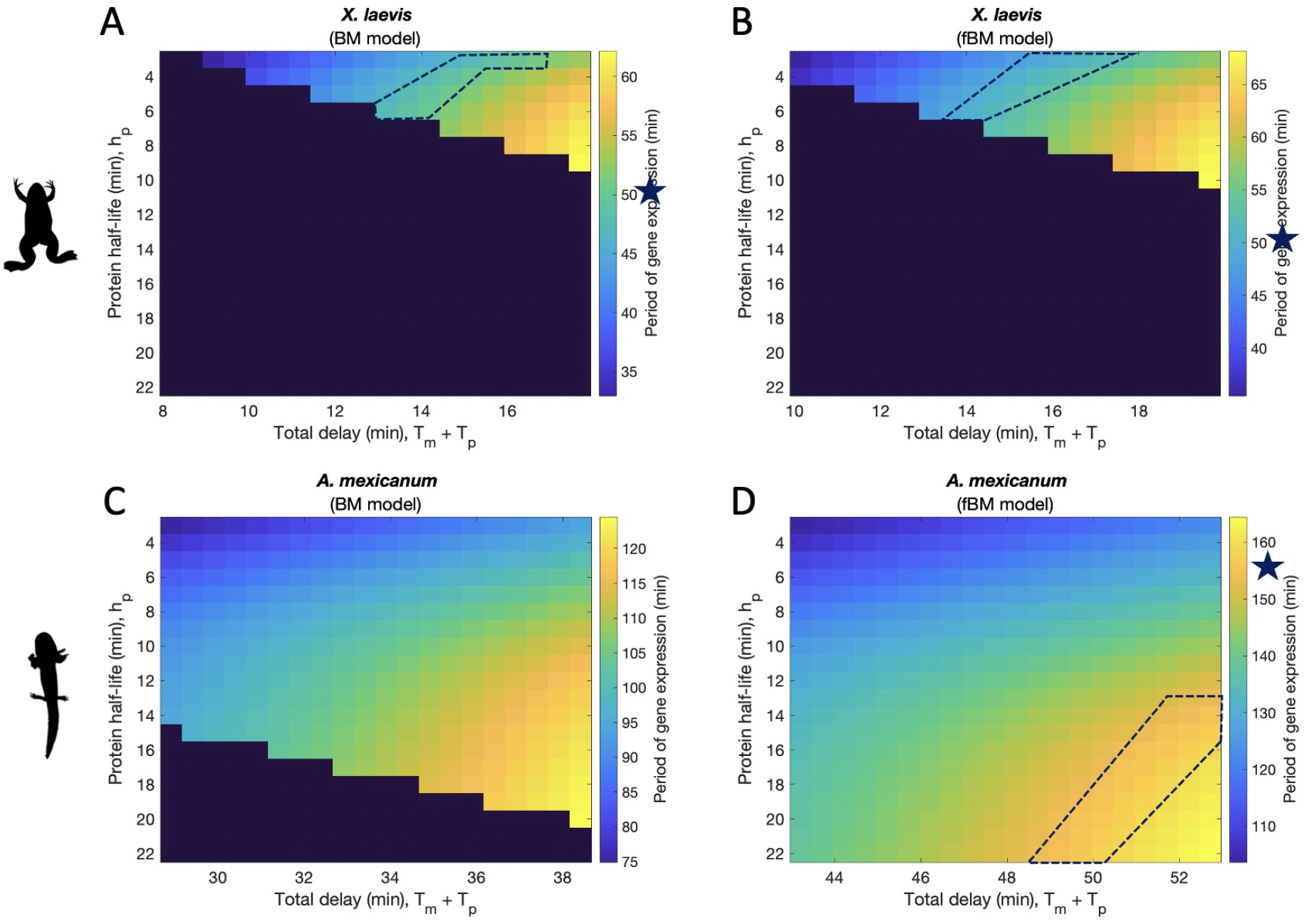
Resulting periods of gene expression given for each species- and diffusion-specific model. Each combination of protein half-life and total delay time corresponds to a period shown in the colorbar to the right of each model plot. Dark purple areas are non-oscillatory; stars show where the known species-specific rates of somite segmentation are found on the colorbar, and the corresponding regions are outlined in dashed lines. Results for: A *X. laevis* model when normal diffusion is assumed/Brownian Motion is modeled; B *X. laevis* model when obstructed diffusion is assumed/fractional Brownian Motion is modeled; C *A. mexicanum* model when normal diffusion is assumed/Brownian Motion is modeled; D *A. mexicanum* model when obstructed diffusion is assumed/fractional Brownian Motion is modeled.

### mRNA export is 2- to 3-fold slower in A. mexicanum relative to X. laevis

mRNA export time roughly doubles in *A. mexicanum* relative to *X. laevis* when normal diffusion is assumed, and roughly triples when obstructed diffusion is assumed (see *T_exp_* in Table 2). The impact of obstructed diffusion on export time becomes more pronounced (i.e. the gap between export times for the two diffusion types becomes wider) as nuclear radius and therefore volume increase (see figure 3).

### Scaling mRNA stability with nuclear export time yields biologically plausible recapitulation of slowed developmental tempo

Although the fBM model shown in figure 1C yields the known rate of somite segmentation in *A. mexicanum* based solely on genome and cell size differences in parameter values, an additional increase in mRNA stability seems logically necessary given our simulated nuclear export times.

The models shown in figure 1 assume an mRNA degradation rate associated with a half-life of 3 minutes. However, we are working with simulated mean export times of *∼*12 and *∼*26 minutes in *A. mexicanum* PSM nuclei under normal and sub-diffusive conditions, respectively. Under these assumptions, a vast majority of mRNA molecules are expected to degrade long before ever leaving the nucleus.

We therefore test if the *A. mexicanum* segmentation clock can also be recovered when mRNA stability is increased relative to export time such that a greater proportion of transcripts are able to exit the nucleus before degrading. Although it is established that some fraction of RNA transcripts will degrade in the nucleus before exiting into the cytoplasm, the extent of degradation seems to differ across transcript types and remains relatively understudied (Smalec et al. 2022). Our parameter estimates for *X. laevis* yield the following patterns: under normal diffusion, mRNA half-life (3 minutes) is *∼*47% of mRNA export delay (6.32 minutes), and under obstructed diffusion, mRNA half-life (3 minutes) is *∼*36% of mRNA export delay (8.41 minutes). In terms of degradation, *∼*76% of mRNA transcripts would be expected to degrade before leaving the nucleus (*∼*24% would leave the nucleus before degrading) under normal diffusion. Under obstructed diffusion, *∼*82% of mRNA transcripts would be expected to degrade before leaving the nucleus (*∼*18% would leave the nucleus before degrading). Meanwhile, when we compare the typical estimate for mRNA half-life in *Danio rerio*, also 3 minutes, to the reported in vivo export time, 3.36 minutes (Hoyle and Ish-Horowicz 2013), we have that mRNA half-life is 83% of export time. This corresponds to an expectation that *∼*58% of mRNA transcripts would degrade before leaving the nucleus (*∼*42% would leave before degrading).

To account for the wide range of empirical and theoretical estimates of mRNA degradation in the nucleus, we explore 3 potential scaling scenarios for mRNA stability and nuclear export time in *A. mexicanum*. We first set mRNA half-life equal to diffusion-specific export times, *h_m_* = *T_exp_*, for which 50% of mRNA transcripts degrade before leaving the nucleus (and 50% leave the nucleus before degrading); then we set mRNA half-life equal to 50% of the simulated export times, *h_m_* = 1/2 *T_exp_*, for which 75% of mRNA transcripts degrade before leaving the nucleus (and 25% leave before degrading); finally, we set mRNA half-life equal to 25% of the simulated export times, *h_m_* = 1/2 *T_exp_*, for which 87.5% of mRNA transcripts degrade before leaving the nucleus (and 12.5% leave before degrading).

In Figure 2, we show plots of *A. mexicanum* model results under both normal and fractional BM conditions, while decreasing the level of mRNA stability. When normal diffusion of transcripts through the nucleoplasm is assumed, figures 2A, 2C, and 2E, none of the mRNA stability scenarios produce a subset of protein half-life and total delay time parameter values that yield a period of 155 minutes. Under obstructed diffusion, figures 2B, 2D, and 2F, all mRNA stability scenarios produce a subset of protein half-life and total delay combinations that fully yield a period of *∼* 155 minutes, matching the known rate of somite segmentation in *A. mexicanum*.

**Figure 2:**
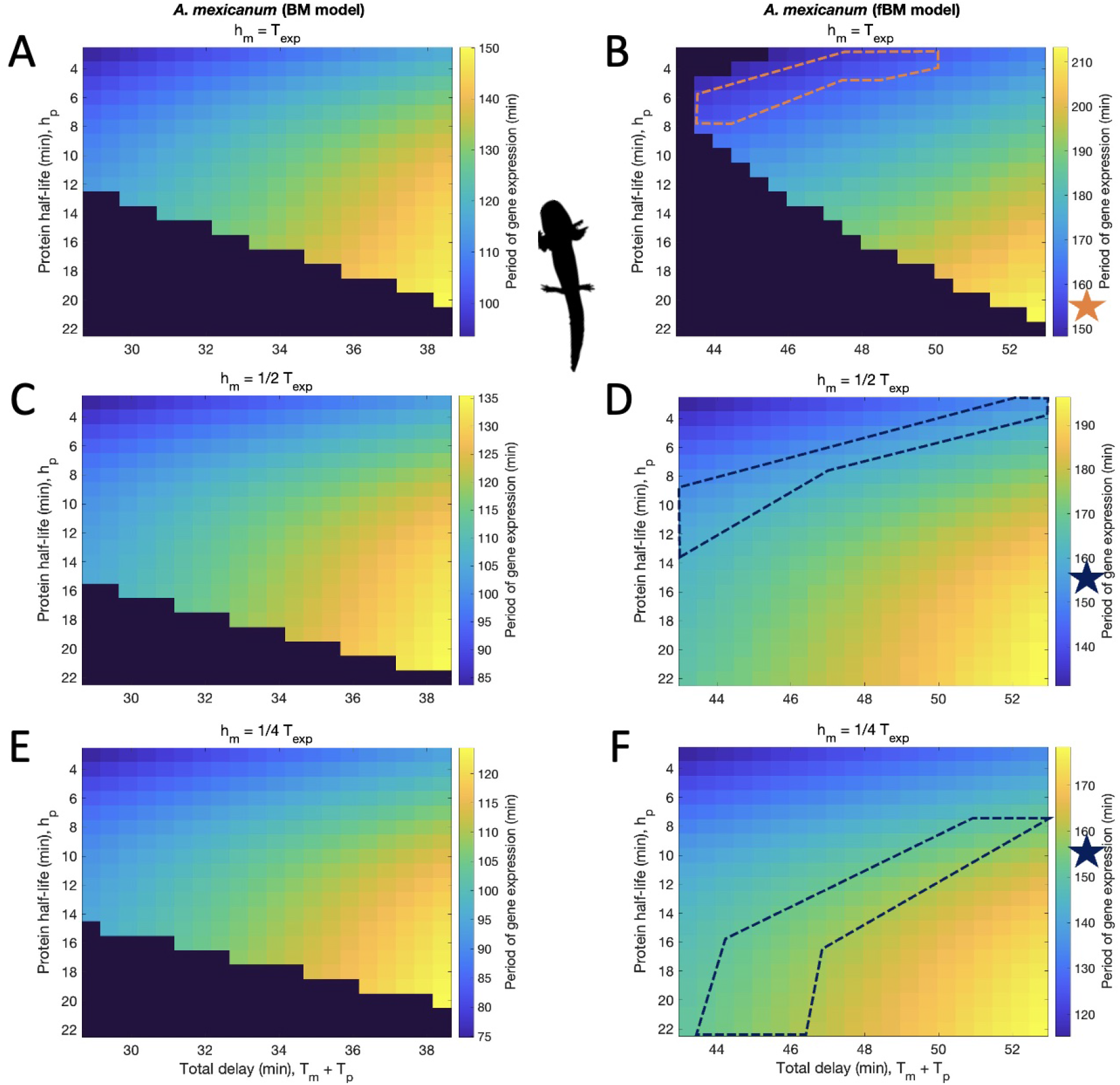
Resulting periods of gene expression for *A. mexicanum* models: A, C, E normal diffusion/Brownian Motion; B, D, F obstructed diffusion/fractional Brownian Motion. mRNA half-life is held constant at: A, B diffusion-specific estimates for mRNA export delay; C, D half of estimated mRNA export delays; E, F a quarter of estimated mRNA export delays. ∗ Note: Color of star/outline is chosen for contrast and has no additional meaning.

## Discussion

Previous research on developmental timing has pointed towards intron length and differences in biochemical characteristics, like degradation rates and network delays, as sources of species-specific tempo (Lázaro et al. 2023, Matsuda et al. 2020, Rayon et al. 2020, Swinburne and Silver 2008, Swinburne et al. 2008). In our study, we not only reconsider these points while contextualizing them within the scope of increasing genome and cell size, but we also make spatially explicit considerations that set our study apart from others.

In Table 2, we outline how species-specific parameters differ between *X. laevis* and *A. mexicanum*, and we note whether each change (based on our methods of estimation) captures increases in genome size or nuclear volume, or neither. Using these species- and diffusion-specific parameters, we first verified that our parameter estimates for *X. laevis* were able to recapitulate the *∼*50 minute segmentation clock. This indicated that our methods of parameter estimation were reasonable, and we moved forward using the same methods for our *A. mexicanum* models, testing for parameter changes and combinations that recapitulate the *∼*155 minute segmentation clock. The results of our models reveal the following potential mechanisms through which developmental tempo slows with increased genome and nuclear size.

### Increasing intron length impacts developmental tempo through transcriptional delays

Increasing the sum of delays in the gene regulatory network has the most significant impact on the period of gene expression (see Table 3: Sensitivity Analysis). When we break total delay down into individual components, we see that while transcriptional delay, *T_tx_*, is not the most significant contributor to total delay, *T_m_* + *T_p_*, it confirms an intuitive link between increasing genome size and the alteration of oscillatory gene expression kinetics. This is because transcriptional delay includes the time needed to transcribe the entire primary gene sequence, including intronic regions, which scale positively with genome size. Indeed, increasing intron size (in part due to transposable element insertions) is a major driver of genomic expansion in *A. mexicanum* (Nowoshilow et al. 2018), and while intronic regions only constitute *∼*17% of the *hes5.7L* primary sequence in *X. laevis* (279 bp of intronic sequence), they make up *∼*76% of the *hes7* primary sequence in *A. mexicanum* (6,307 bp of intronic sequence). This is a general pattern observed across orthologous genes in *A. mexicanum* and *X. laevis* (Nowoshilow et al. 2018). Interestingly, this pattern of intronic expansion seems to be constrained in developmental genes, where an *∼*11-fold increase in average intron length is observed between *X. laevis* and *A. mexicanum* in contrast with an almost 20-fold increase seen in non-developmental genes (Nowoshilow et al. 2018). In Nowoshilow et al. [2018], the role of transcription in developmental timing is suggested as an explanation for this pattern. The time required to transcribe intronic regions of genes (i.e. “intron delay”) has not only been identified as a potential mediator of developmental timing via its role in gene regulatory networks (Swinburne and Silver 2008), but has also been suggested to link genomic gigantism in particular with slowed development and regenerative abilities through its impact on gene expression kinetics (Sessions and Wake 2021).

**Table 3:** Sensitivity analysis.

| Original parameter set | Period of mRNA expression | Amplitude of mRNA expression |  |  |
| --- | --- | --- | --- | --- |
|  | (minutes) | (molecules) |  |  |
| $a = 4.5, k = 33, p_{crit} = 420,$ | | | | |
| $h_m = 3, h_p = 15,$ | | | | |
| Total delay = 33.91 | 109.08 |  | 31.21 |  |
| (A. mexicanum BM model) |  |  |  |  |
| Parameter changed | 50% increase | 50% decrease | 50% increase | 50% decrease |
| $a$ | 110.02 | 107.84 | 28.76 | 26.23 |
| $k$ | 109.93 | 107.87 | 43.13 | 13.01 |
| $p_{crit}$ | 108.41 | 110.48 | 30.58 | 26.19 |
| $h_m$ | 114.55 | 102.88 | 41.84 | 11.77 |
| $h_p$ | Inf | 98.41 | 0.99 | 112.07 |
| Total delay ($T_m + T_p$) | 152.79 | Inf | 78.34 | 0.05 |

However, according to our computational results, increased transcriptional delay alone cannot drive the vast differences in overall delay time between *X. laevis* and *A. mexicanum*. Our model therefore also implies that intron delay alone cannot drive slowed developmental tempo observed in the *A. mexicanum* segmentation clock relative to that of *X. laevis*. Instead, we see in Table 2 that the delay associated with increasing nuclear volume, *T_exp_* is the most significant contributor to overall delay time across all species- and diffusion-specific models. Although the delay associated with intron-splicing, *T_in_* is an important contributor to total delay, and the difference between species-specific values for *T_in_* is in fact larger than that between species-specific values for *T_tx_*, inter-species differences in splicing kinetics cannot be clearly linked with differences in intron or, by extension, genome size (Khodor et al. 2012).

### mRNA export time increases with nuclear volume and has a pronounced impact on developmental tempo

Increasing nuclear volume between *X. laevis* and *A. mexicanum* is captured by the critical protein threshold value, *p_crit_*, and mRNA export delay, *T_exp_*, parameters. The increase in critical protein threshold accounts for the fact that in a larger nucleus, more protein molecules are needed to reach the same critical concentration of 10*^−^*^9^ *M* required for transcriptional repression to act “in earnest” (Lewis 2003). However, changes to *p_crit_* have a very limited impact on the period of gene expression relative to other parameter changes. Meanwhile, the increase in mRNA export delay in *A. mexicanum* relative to *X. laevis* has a relatively pronounced impact on the period of gene expression.

Although the increase in the estimated radius of PSM nuclei in *A. mexicanum* relative to *X. laevis* may seem minimal, 5.5*µ*m compared to 4*µ*m, mRNA export delay still nearly doubles in *A. mexicanum* when normal diffusion is assumed and nearly triples when obstructed diffusion is assumed. The impact of increasing radius is greater when obstructed diffusion is assumed, and differences in estimated export times (between lengths and diffusion types) become more pronounced as radius increases. As a result, we would expect mRNA export delays to become more pronounced in gene regulatory networks across increasing nuclear volume, and we might also expect mRNA export to be the largest contributor to total delay in a gene regulatory network, especially in larger nuclei. This can be seen in figure 3, where we plot simulated mean export times under both normal and obstructed diffusion for radii between 3 and 6 *µ*m. If we expand the range of radii past 6 *µ*m, the impact of increasing radius and obstructed diffusion on export time becomes even more pronounced, suggesting that the nuclear center in species with the largest known genome sizes may have become “uninhabitable” for genes with dynamic expression patterns. If we introduce transcripts whose positions are drawn from a uniform distribution within the nucleus, as opposed to always starting from the origin/nuclear center, the impacts of increasing radius and obstructed diffusion on mean export time are reduced. The distribution of export times becomes more skewed towards lower values (because most transcript trajectories are starting closer to the periphery) (see Supplemental Material 3). In very large nuclei, we hypothesize that genes – especially those with dynamic expression patterns – may be constrained to occupy these locations away from the nuclear center, suggesting an overall effect of genome expansion on the organization of chromosomal territories.

**Figure 3:**
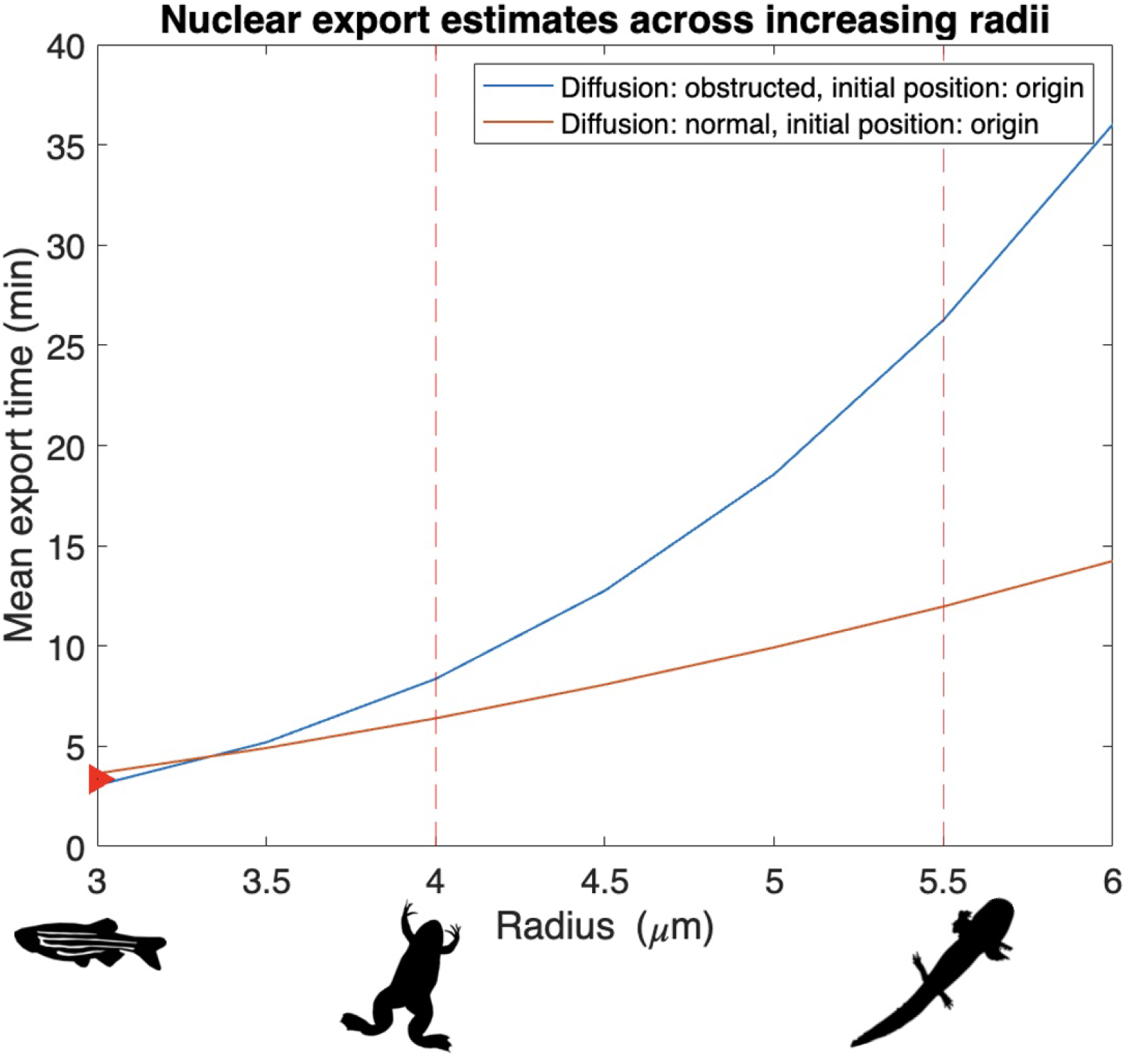
Nuclear export estimates under normal and obstructed diffusion across a range of radii that captures estimates for *X. laevis* and *A. mexicanum* PSM nuclei (shown by the red dashed lines). An initial position at the nuclear center is assumed. Trajectories are scaled such that a radius of 3*µ*m corresponds to a mean export time of *∼*3.36 minutes (shown by the red arrow) to match the reported export time of *her1 (hes7)* in zebrafish Hoyle and Ish-Horowicz 2013. ∗ Note: There is a limit to how closely 3*µ*m can be scaled with *∼*3.36 minutes. As a result, obstructed diffusion mean export times start off slightly faster than normal diffusion when radius *r <* 3.5*µ*m, but this is not necessarily biologically meaningful, and mean export times are quick to converge back to expectations.

Regardless of model type, assuming an intial position at the nuclear center, our simulations position mRNA export as the largest contributor to total delay time. This is consistent with observations that find mRNA export delays to be longer than both transcriptional and intron splicing delays in mice, chick, and zebrafish segmentation clocks (Hoyle and Ish-Horowicz 2013). As the largest contributor to total delay time, mRNA export is also the largest contributor to overall differences in *X. laevis* and *A. mexicanum* segmentation clocks, implying that spatially induced delays in gene regulatory networks play a significant role in the slowing of cellular and developmental processes. This suggests an important albeit underexplored spatial component in the mediation of developmental tempo across increasing cell and nuclear volumes.

### Simulations position increasing chromatin density as a driver of slowed development

Across the tree of life, decreasing nucleus to cell volume ratios are observed as cells increase in volume (Malerba and Marshall 2021). In other words, larger cells have relatively smaller nuclei, though absolute nuclear volumes still increase with cell volume. Given the well-established positive correlation between genome size and cell volume, we might also state that nuclei become relatively smaller with increasing genome size. As a result, we would see an increasing amount of genetic material packed within relatively smaller nuclei, implying greater chromatin packing density as genome size and absolute nuclear volumes increase. Readers will recall that chromatin structure density is a commonly cited source of obstructed diffusion observed in the nucleus Ben-Ari et al. 2010, Ishihama and Funatsu 2009, Mor and Shav-Tal 2010, Oeffinger and Zenklusen 2012, Sheinberger and Shav-Tal 2013, meaning that we would expect to see obstructed diffusion of transcripts in absolutely larger nuclei/organisms with relatively large genomes.

With these empirical possibilities in mind, we turn towards our theoretical results. In figures 1A and 1B, we see that both diffusion models are able to fully recapitulate the *X. laevis* segmentation clock, while obstructed diffusion (modeled by fractional Brownian Motion, shown in figure 1D) must be assumed to fully recapitulate the slowed *A. mexicanum* segmentation clock. Similarly, in figure 2, we have that obstructed diffusion still must be assumed, regardless of mRNA stability, in order to recapitulate the slowed *A. mexicanum* segmentation clock. Meanwhile, under normal Brownian Motion, all model scenarios fall short of fully recapitulating the *A. mexicanum* segmentation clock. Further increases in protein and/or mRNA stability do not fully resolve this issue (see Supplemental Material 4), suggesting that normal diffusion does not introduce sufficient delays into the system. These results indicate that obstructed diffusion is sufficient but not necessary to recapitulate the segmentation clock when genome size is relatively small (*X. laevis*) yet becomes both sufficient and necessary when genome size is relatively large (*A. mexicanum*), supporting the hypothesis that chromatin packing density increases with genome size. Somitogenesis transcripts must exhibit obstructed diffusion in the nucleoplasm of *A. mexicanum* in order to slow the segmentation clock to an appropriate pace, thus positioning increasing chromatin density as a potential driver of slowed development.

### Gene product stability also acts to mediate developmental tempo

Focusing on figures 2B, 2D, and 2F (fBM model results), we can see that as mRNA stability changes, so too does the range of protein half-life and total delay time combinations yielding a gene expression period of 155 minutes. This is in agreement with previous studies that point to gene product stability as a mediator of species-specific segmentation clock periods (Hirata et al. 2004, Lázaro et al. 2023, Matsuda et al. 2020, Rayon et al. 2020). However, some mRNA stability scenarios yield more reasonable results than others. In figures 2B and 2D, with mRNA half-lives of 26.27 and 13.14 minutes, the 155 minute period of oscillation is captured for protein half-lives that range from 3 to 8 and 3 to 13 minutes, respectively. As a result, we would have to assume that protein is as stable or less stable than mRNA, which is inconsistent with reported patterns for Hes7 in mice and humans (Matsuda et al. 2020). In figure 2F, for which mRNA half-life is set to 6.54 minutes, we get a larger subset of parameter combinations that yield a period of 155 minutes. Across this subset, protein half-lives range from being only slightly more stable than mRNA at *∼*8 minutes to almost 4-times as stable as mRNA at *∼*23 minutes. The resulting relationship between mRNA and protein stability, that protein molecules exhibit increased stability relative to their transcripts, is more consistent with the patterns reported in Matsuda et al. [2020]. However, setting 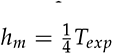 implies that a vast majority of mRNA transcripts, 87.5%, will degrade before exiting the nucleus. We might expect amplitude (i.e. number of transcripts in the cell) to be extremely low as a result, but this is not the case. In fact, mRNA degradation is related in complicated ways to not only the amplitude of mRNA expression, but to the amplitude of protein expression as well, resulting in counter-intuitive patterns of expression across different mRNA stability scenarios (see Supplemental Material 5).

Considering again the period of gene expression, an interesting tradeoff exists between the possible mRNA stability scenarios described above: in order to mathematically recapitulate the slowed *A. mexicanum* segmentation clock, we must either assume that a vast majority of mRNA transcripts degrade within the nucleus, or that the protein molecules in this system are less stable relative to their transcripts. The assumption of obstructed diffusion necessitates some kind of deviation – either high rates of nucleoplasmic degradation, or low stability of protein molecules relative to their transcripts – from conventional gene expression kinetics as genome and nuclear size increase between *X. laevis* and *A. mexicanum*. Based on reported patterns for Hes7 in mice and humans (Matsuda et al. 2020), we propose that widespread nucleoplasmic degradation of the somitogenesis transcript is more plausible than transcripts that are more stable than the corresponding protein molecules.

What remains the same across genome size and nuclear volume is that, as shown in previous implementations of the Lewis model (Hirata et al. 2004, Matsuda et al. 2020), the relative stabilities of different gene products interact to shape clock tempo. This is demonstrated by changing ranges of protein half-life corresponding to a period of 155 minutes as mRNA half-life decreases. However, gene product stability alone cannot capture slowed development across increasing genome size and nuclear volume. Simply increasing protein stability and/or mRNA stability for the Brownian Motion (normal diffusion) models, shown in figures 2A, 2C, and 2E, cannot fully recapitulate the slowed *A. mexicanum* clock (see Supplemental Material 4 for an extended discussion). The additional delays introduced by obstructed nucleoplasmic diffusion are necessary to sufficiently slow the period of gene expression. Although gene product stability acts to mediate developmental tempo, our model results suggest that across increasing genome size and nuclear volume, gene product stability does not act alone to slow developmental tempo.

## Conclusion

Our results suggest that spatial differences must be accounted for to recapitulate slowed development as a result of evolutionary increases in genome and cell size. In agreement with previous studies, our results also suggest an important role for gene product stability when it comes to mediating species-specific rates of development. Taken together, we show that the physical and spatial delays predicted by increased intron length and nuclear size, coupled with alterations to gene product stability that ensure the products persist for long enough to fulfill their molecular function, mathematically recapitulate the slow developmental tempo found in species with large genomes.

## Data and Code Availability

Simulation code and data can be accessed at: https://github.com/ataylor3/Amphibian-segmentation-clock-models

