## Supplemental Material for "Amphibian segmentation clock models suggest mechanisms of slowed development across increasing genome size and nuclear volume"

*Manuscript elements*: Table S1, Figure S1, Figure S2, Figure S3, Figure S4, Figure S5, Figure S6, Figure S7

Prepared using the suggested LaTeX template for *Am. Nat.*

### Supplement 1: Differences between periods of mRNA and protein expression are minimal

Table S1: Resulting periods of mRNA and protein expression given parameter values corresponding to *X. laevis* and *A. mexicanum* Brownian Motion models

| Parameter set | **Resulting period of mRNA expression** (minutes) | **Resulting period of protein expression** (minutes) |
|---|---|---|
| $a = 4.5$, $k = 33$, $p_{crit} = 161$, $h_m = 3$, $h_p = 3$, Total delay = 13.17 (*X. laevis* BM model) | 43.03 | 43.04 |
| $a = 4.5$, $k = 33$, $p_{crit} = 420$, $h_m = 3$, $h_p = 3$, Total delay = 33.91 (*A. mexicanum* BM model) | 84.97 | 85.03 |

### Supplement 2: Analytical conditions for the emergence of oscillations

Following the derivation given in Verdugo and Rand [2008], we derive formulae that give analytical conditions for the emergence of oscillations for the system of delayed differential equations (DDE) modelling the segmentation clock as proposed by Lewis [2003].

We begin with Lewis' DDE system, using $\dot{p}$ and $\dot{m}$ derivative notation in place of $\frac{dp}{dt}$ and $\frac{dm}{dt}$, respectively, for convenience:

$$\dot{p} = am(t - T_p) - bp(t) \tag{1}$$

$$\dot{m} = \frac{k}{1 + (\frac{p(t-T_m)}{p_{crit}})^2} - cm(t) \tag{2}$$

and we start by rescaling the system.

**Rescaling** We rescale, following Verdugo and Rand, and define the following variables:

$$x = \frac{m}{k}, \quad y = \frac{p}{ka}, \quad Y_0 = \frac{p_{crit}}{ka} \tag{3}$$

We also use the following notation: $m_d = m(t - T_p)$ and $p_d = p(t - T_m)$, and we define $x_d$ and $y_d$ similarly. From the variables defined in Equation (3), we have:

$$\dot{x} = \frac{1}{1 + (y_d/Y_0)^2} - cx, \tag{4}$$

$$\dot{y} = x_d - by \tag{5}$$

**Steady state solution** At the steady state $(x^*, y^*)$ we have $x_d = x^*$ and $y_d = y^*$. Setting $\dot{x} = 0$ and $\dot{y} = 0$ yields the following steady state equations:

$$\frac{1}{1 + (y^*/Y_0^2)} - cx^* = 0 \tag{6}$$

$$x^* - by^* = 0 \tag{7}$$

Eqn. (7) implies $x^* = by^*$. Upon substituting in Eqn. (8) we get the following cubic equation for $y^*$:

$$(y^*)^3 + Y_0^2(y^*) - Y_0^2/cb = 0 \tag{8}$$

This equation can be solved by Mathematica. It has one real root given by,

$$y^* = -\frac{2^{1/3}Y_0^2}{D} + \frac{D}{3 * 2^{1/3}} \tag{9}$$

where,

$$D = \left[\frac{27Y_0^2}{bc} + 3\sqrt{\frac{81Y_0^4}{b^2c^2} + 12Y_0^6}\right]^{1/3} \tag{10}$$

**Linearize about a fixed point** To linearize about a fixed point we define the following deviations from the steady state $(x^*, y^*)$,

$$\xi = x(t) - x^*, \quad \eta = y(t) - y^* \tag{11}$$

and we use the subscript d to signify the lagged variable such that $\xi_d = \xi(t - T_m)$ and $\eta_d = \eta(t - T_p)$. Note also that $\xi_d = x(t - T_m) - x^*$ and $\eta_d = y(t - T_p) - y^*$.

It follows that,

$$\dot{\xi} = \dot{x} = \frac{1}{1 + \frac{(\eta_d + y^*)^2}{Y_0^2}} - c(\xi + x^*) \tag{12}$$

and,

$$\dot{\eta} = \dot{y} = \xi_d + b\eta \tag{13}$$

We now expand Eqn. 12 for small $\eta_d$. To linear order we get,

$$\dot{\xi} = -c\xi - cx^* + \frac{Y_0^2}{(y^*)^2 + Y_0^2} - \frac{2y^*(Y_0)^2}{((y^*)^2 + Y_0^2)^2} * \eta_d \tag{14}$$

Note that the second and third terms on the right hand side of Eqn. (14) sum to zero.

In order to compare the results in Eqn. (14) to Verdogu and Rand, we rewrite the coefficient of $\eta_d$ in terms of $\beta = y^*/Y_0$ and define:

$$K = \frac{2y^*(Y_0)^2}{((y^*)^2 + Y_0^2)^2} = \frac{2\beta^2}{y^*(1 + \beta^2)^2} \tag{15}$$

We therefore end up with the following linearized equations:

$$\dot{\xi} = -x\xi - K\eta_d \tag{16}$$

$$\dot{\eta} = \xi_d - b\eta \tag{17}$$

which are analogous to those presented in Verdugo and Rand

**Oscillatory solution** In a supercritical Hopf bifurication, a stable fixed point becomes unstable and is surrounded by a stable limit cycle. Close to the bifurcation, the amplitude of the limit cycle is very small and can be approximated by cosine functions. Therefore, we assume that Eqns. 16 and 17 have solutions given by,

$$\xi(t) = Bcos(\omega t + \phi), \xi_d = Bcos(\omega(t - T_p) + \phi) \tag{18}$$

$$\eta(t) = Acos(\omega t), \eta_d = Acos(\omega(t - T_m)) \tag{19}$$

We can now substitute Eqns. (18) and (19) into the linearized equations, Eqns. (16) and (17), above to solve for $\omega$ in terms of model parameters. The condition for oscillation is thus that $\omega$ is a real, positive, non-zero number. After some tedious trigonometry and algebra, we find that this condition is satisfied when $K > bc$. In other words the geometric mean of the degradation constants has an upper bound given by the following.

$$\sqrt{bc} < \sqrt{K} \tag{20}$$

The frequency of oscillation $\omega$ is given by:

$$\omega = \left[ \frac{-(c^2 + b^2) + \sqrt{(c^2 + b^2)^2 + 4K^2 - 4b^2c^2}}{2} \right]^{1/2} \tag{21}$$

We may also ask if there exists a minimum total delay, $T = T_p + T_m$, required for the emergence of oscillations. Solving for such a $T$, we get:

$$T = \frac{1}{\omega} ArcSin \left[ \frac{\omega(c + b)}{K} \right] = T_{crit} \tag{22}$$

and we define this $T$ to be a critical total delay, $T_{crit}$, because it is the time delay required for an oscillatory solution just at the bifurcation. It follows that a second condition for oscillation can be defined, that is:

$$T_m + T_p > T_{crit} \tag{23}$$

Note that even when the conditions given in (20) and (23) are met, a finite cut-off in numerical simulations may lead a solution to be classified as non-oscillatory.

In figures S1 and S2, we plot $h_p$ against $T_{crit}$ for every model shown in figures 1 and 2 in the main text, respectively (note that the axes are swapped relative to their corresponding plots in the main text, to clearly demonstrate the positive relationship between protein half-life and critical total delay). $T_{crit}$ is dependent on protein and mRNA degradation rates, $b$ and $c$, respectively, as well as $K$. $K$ is dependent on $\beta$ and $y^*$, both of which are dependent on $Y_0$ which is defined as $Y_0 = p_{crit}/(ka)$. As a result, we can say that $T_{crit}$ is dependent on only three parameters from the Lewis model: $b$, $c$, and $p_{crit}$. Therefore, we need only generate multiple species-specific plots when $h_m$ is changing between models (recall that $b = ln(2)/h_p$ and $c = ln(2)/h_m$) because the range of $h_p$ and value of $p_{crit}$ will remain the same across all species-specific models, regardless of diffusion type. This is why there are only two plots in figure S1 (corresponding to species- and diffusion-specific for which mRNA half-life is held constant at $h_m = 3$), but there are six plots in figure S2 (corresponding to diffusion-specific *A. mexicanum* models for which $h_m$ is adjusted to different scaling relationships with diffusion-specific $t_{exp}$ values).

There is a clear and consistent trend across all plots in both figures. That is, as protein half-life increases, the total delay time required for oscillations to emerge, $T_{crit}$, also increases. Analytical analysis therefore confirms that total delay time must increase with protein stability to ensure the emergence of oscillations

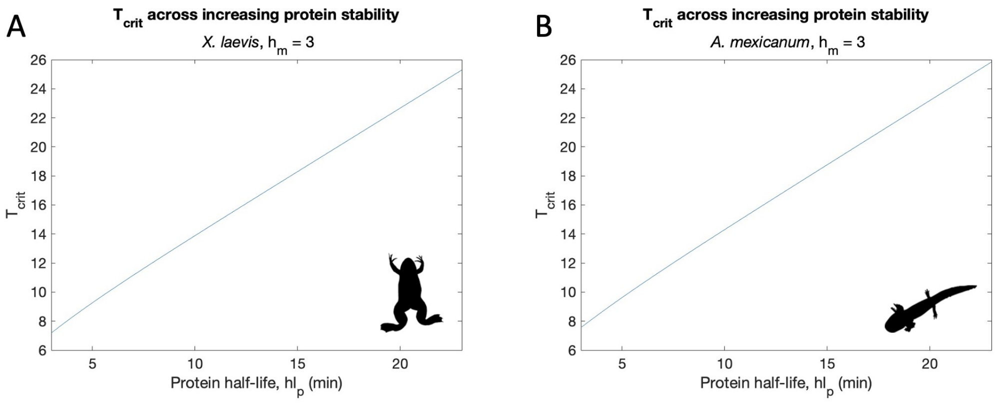


Figure S1: $T_{crit}$ plotted across increasing protein stability corresponding to a range of half-lives between 3 and 23 minutes. A *X. laevis* BM and fBM models for which $h_m = 3$, as shown in figures 1A and 1B. B *A. mexicanum* BM and fBM models for which $h_m = 3$, as shown in figures 1C and 1D.

### Supplement 3: Extended nuclear export simulation results

Extending nuclear export simulations across a larger range of radii reveals that the impact of obstructed diffusion is pronounced in larger nuclei, but it is reduced by drawing initial positions from a uniform distribution.

In figure S3A we show nuclear export simulation results across a range of interest that captures estimates for nuclear radii of presomitic mesoderm tissue cells in *X. laevis* and *A.mexicanum*, and in figure S3B we show results across a range of ~0.5 to 13 $\mu$m, based on the minimum and maximum nuclear volumes reported in the dataset used by Malerba and Marshall [2021] (`https://doi.org/10.5061/dryad.vq83bk3ss`) while assuming a spherical volume $V = \frac{4}{3}\pi r^3$. It is important to note that while the results in figure S3B give us an idea of how nuclear export delay increases across nuclear radii for different types of diffusion and chromatin addresses, these estimations are still scaled based on observed nuclear export time in zebrafish (Hoyle and Ish-Horowicz 2013), so we cannot make definitive conclusions about nuclear export times for all transcripts in all nuclei of a certain size.

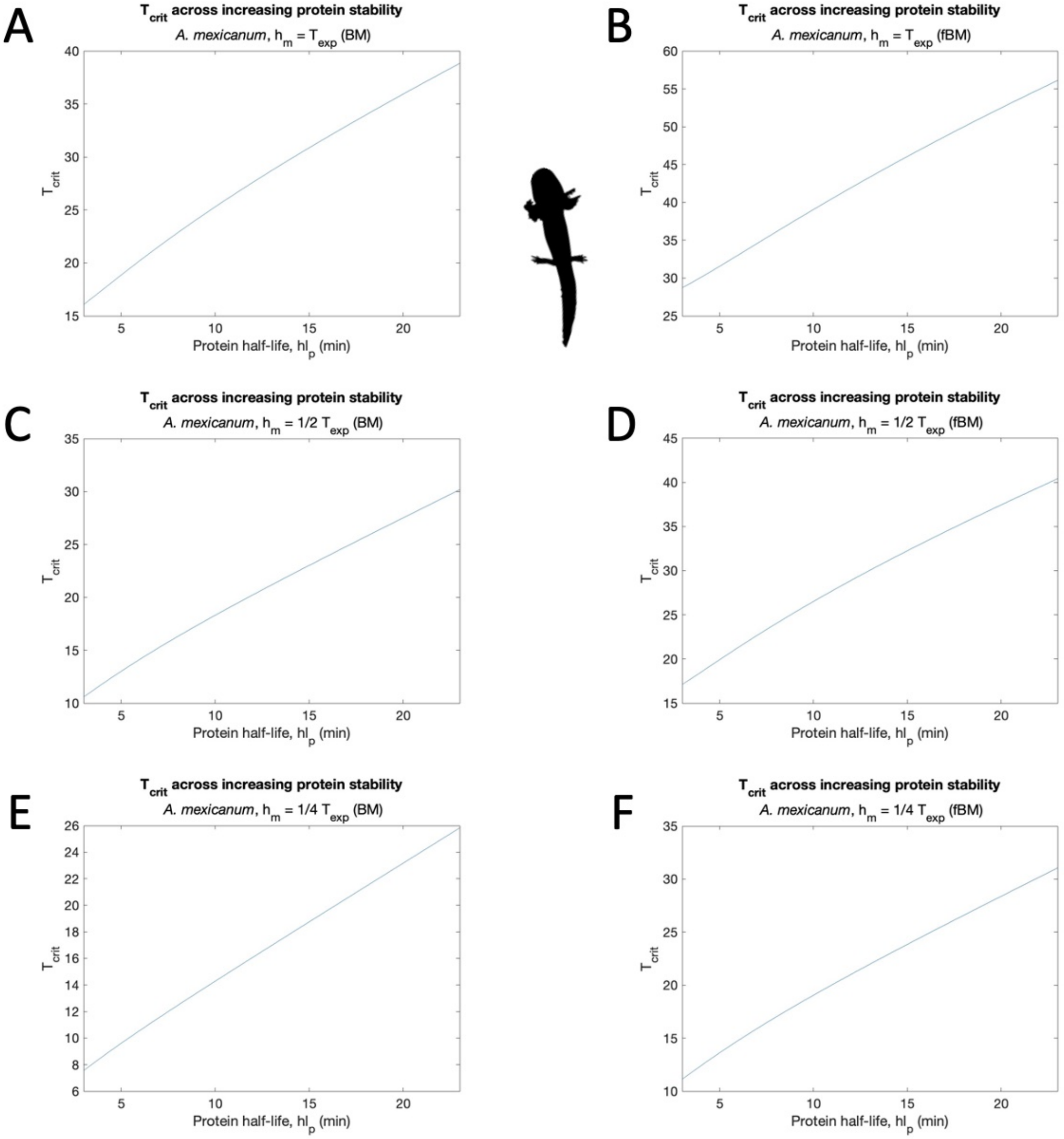


Figure S2: $T_{crit}$ plotted across increasing protein stability corresponding to a range of half-lives between 3 and 23 minutes. A, C, E *A. mexicanum* BM models with $h_m = T_{exp}$, $h_m = \frac{1}{2}T_{exp}$, and $h_m = \frac{1}{4}T_{exp}$, respectively. B, D, F *A. mexicanum* fBM models with $h_m = T_{exp}$, $h_m = \frac{1}{2}T_{exp}$, and $h_m = \frac{1}{4}T_{exp}$, respectively. All panels correspond with their counterparts from figure 2.

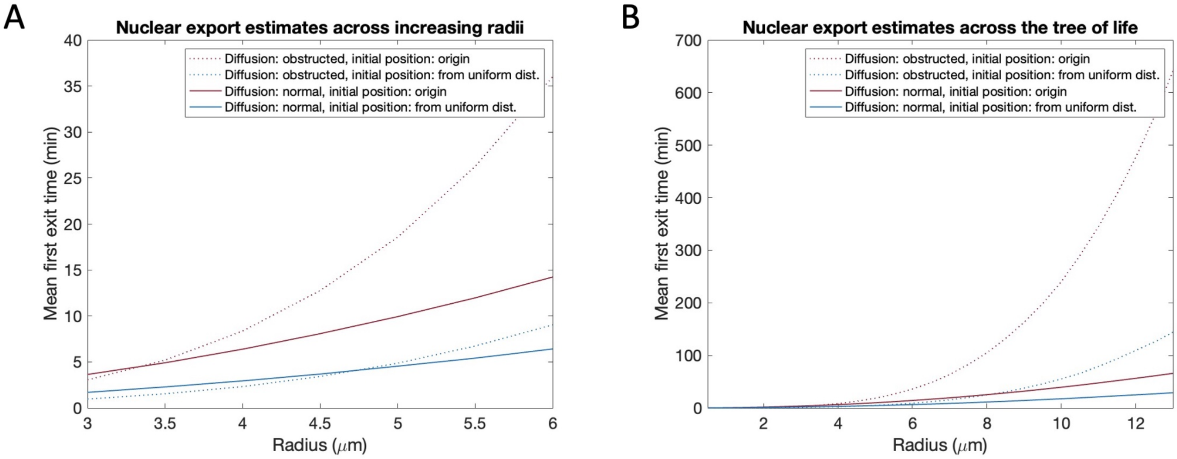


Figure S3: Nuclear export simulation results. A Mean nuclear export time across nuclear radii in *D. rerio*, *X. laevis*, and *A. mexicanum* PSM cells. B Mean nuclear export time across a wider range of nuclei that reflect what has been observed across the tree of life. Simulation results are shown for Brownian Motion(BM)/normal diffusion (solid lines) and fractional Brownian Motion(fBM)/obstructed diffusion (dashed lines), and for initial positions at the origin (shown in maroon) and drawn from a uniform distribution (shown in blue).

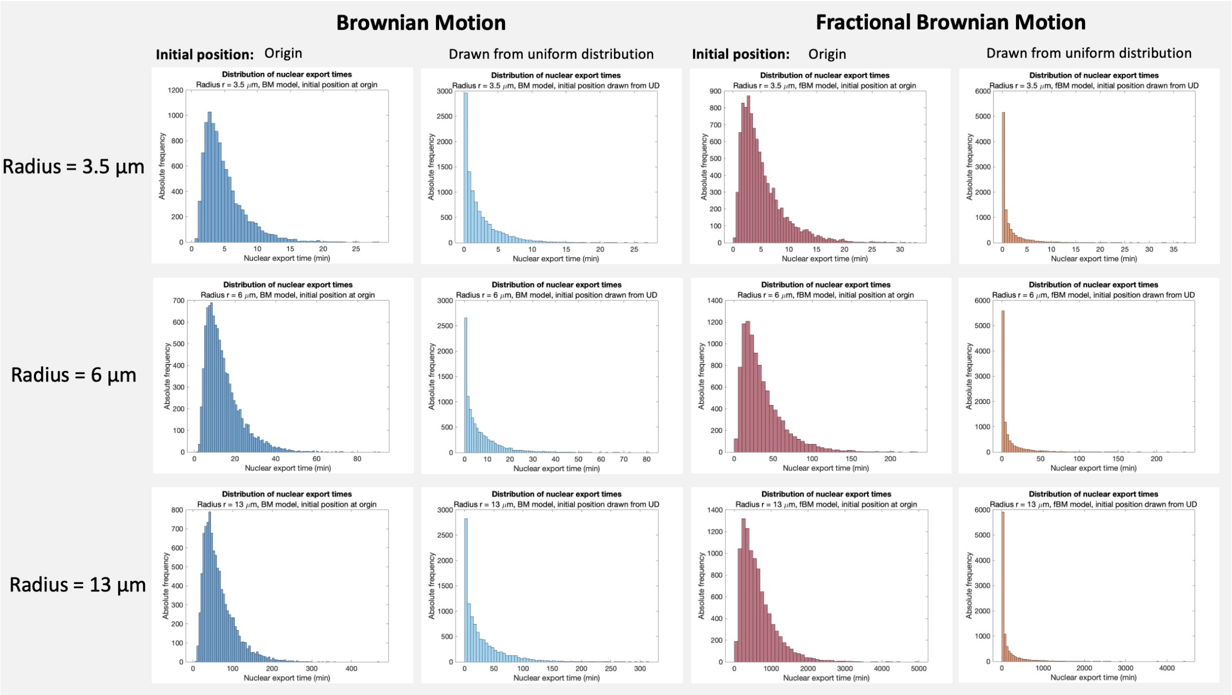


Figure S4: Nuclear export distributions for nuclei with radius 3.5, 6, and 13 $\mu$m, and across different diffusion and initial position models.

In figure S4, we plot the distribution of nuclear export times for all four diffusion models shown in figure S3 (BM with initial position at the origin, BM with initial positions drawn from a uniform distribution, fBM with initial position at the origin, and fBM with initial positions drawn from a uniform distribution), and for three different nuclear radii (3.5, 6, and 13 $\mu$m; most species have nuclear radii well below 6 $\mu$m). We can see that nuclear export distributions are skewed more towards the left (i.e. towards smaller times) when initial positions are drawn from a uniform distribution relative to when initial position is always at the origin or nuclear center. This pattern holds across nuclear size.

### Supplement 4: Increasing gene product stability for *A. mexicanum* Brownian Motion models

Figure 2A corresponds to the Brownian Motion *A. mexicanum* model that is closest to achieving a period of gene expression to match the species-specific segmentation rate of 155 minutes. For this model, we have that $h_m = T_{exp} = 11.97$. Looking at figure 2A, we can imagine that increasing the range of protein stability might yield a period of gene expression slow enough to match the known rate of somite segmentation in *A. mexicanum*, and that the subset of parameters to yield a period of 155 minutes would correspond to protein stability that is relatively high compared to mRNA stability. Furthermore, we would have a scenario in which only 50% of mRNA transcripts are expected to degrade before leaving the nucleus, $h_m = T_{exp}$. To test this, we take the model shown in figure 2A, and we consider a new range of protein stability from 15 to 35 minutes (as opposed to from 3 to 23 minutes). All other parameters are held constant. As shown in figure S5A, solely increasing protein stability while holding all other parameters does not yield the 155 minute rate of somite segmentation.

Increasing mRNA stability by 25%, $h_m = 14.96$, while holding protein stability at its higher range, yields one parameter combination with a period of $\sim$154 minutes (figure S5B). Increasing mRNA stability by 50% relative to $h_m = 17.96$, yields only one parameter combination for which oscillations emerge, and its period is $\sim$153 minutes (figure S5C). We can only begin to recapitulate the *A. mexicanum* segmentation clock (i.e. we are within 2 minutes of the known segmentation rate) under a Brownian Motion/normal diffusion model if both mRNA and protein stability are increased. However, we still fail to fully recapitulate a period of 155 minutes. Without increases in the total delay time, $T_m + T_p$, there is an upper limit on how stable gene products can become before the emergence of oscillations is no longer possible; instead of slowing the segmentation clock down, we "break" it altogether by damping oscillations. Additionally, results in figure S5B and S5C suggest that to achieve a period close to 155 minutes protein molecules are either only slightly more stable than their transcripts (S5B), or less stable than their transcripts (S5C). This is

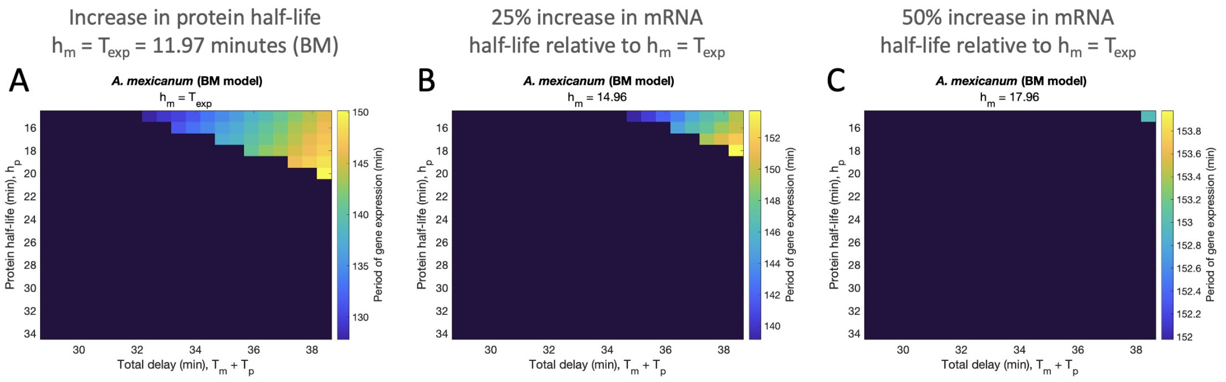


Figure S5: Further increasing gene product stability in the *A. mexicanum* BM model. A We keep mRNA half-life equal to the species- and normal diffusion-specific estimated nuclear export time, and we set protein stability to a higher range corresponding to half-lives of 15 to 35 minutes. B We keep this high range of protein stability constant and we increase mRNA stability by 25% relative to $h_m = T_{exp}$. C We increase mRNA stability by 50% relative to $h_m = T_{exp}$ while holding the high range of protein stability constant.

an issue discussed in the main text.

### Supplement 5: Amplitude plots

In figures S6 and S7, we plot the amplitudes of mRNA and protein expression, respectively, corresponding to the models plotted in figure 2, for which mRNA stability is set to scale with estimated export time, $h_m = T_{exp}$. In figure S6, we can see that when mRNA stability is decreased from $h_m = T_{exp}$ by 50% to $h_m = \frac{1}{2}T_{exp}$, we see an overall increase in the amplitude of mRNA expression, yet when mRNA stability is further decreased to $h_m = \frac{1}{4}T_{exp}$ there is an overall decrease in amplitude. This general pattern is seen across both models (normal and fractional Brownian Motion), and in figure S7, we see this pattern extend to protein as well. The results shown in figures S6 and S7 are counter intuitive. First, we might intuit that higher rates of mRNA degradation (i.e. lower levels of stability) would result in low transcript numbers, but we see the opposite when mRNA stability decreases from $h_m = T_{exp}$ to $h_m = \frac{1}{2}T_{exp}$. One might reason that increased transcript degradation would lead to less translation resulting in lower protein numbers and therefore less transcriptional repression, and that this would explain an increase in transcript numbers following a decrease in mRNA stability. However, we also have increasing protein amplitude when mRNA stability decreases from $h_m = T_{exp}$ to $h_m = \frac{1}{2}T_{exp}$, so this explanation does not hold. Furthermore, the increase in both mRNA and protein amplitude is followed by a decrease when mRNA stability is further decreased from $h_m = \frac{1}{2}T_{exp}$ to $h_m = \frac{1}{4}T_{exp}$. Taken together, we have that there is a complex relationship between mRNA stability and the amplitude of mRNA and protein expression.

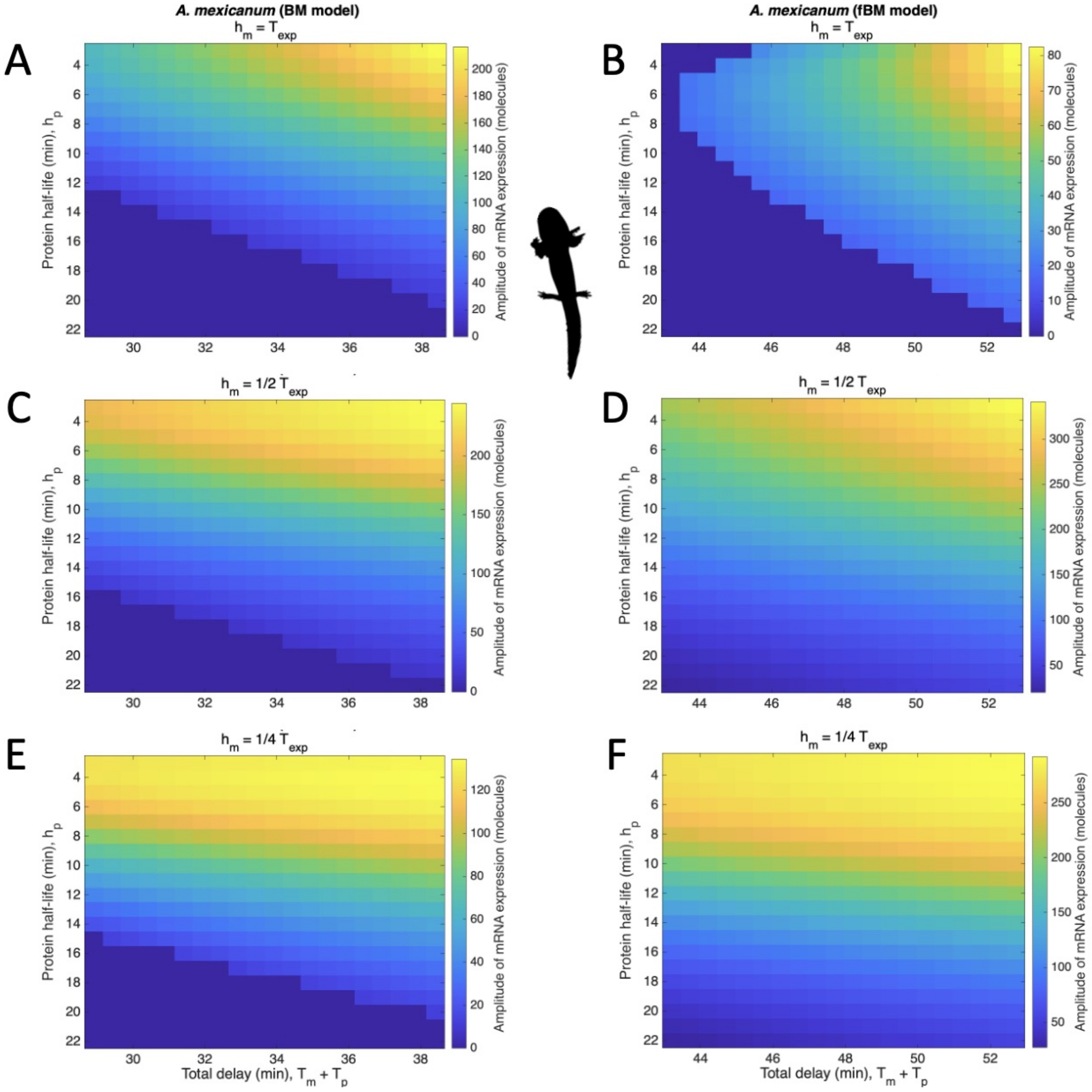


Figure S6: Resulting amplitudes of mRNA expression for *A. mexicanum* models corresponding to plots in figure 2. A, C, E normal diffusion/Brownian Motion model results; C, D, F obstructed diffusion/fractional Brownian Motion model results. mRNA half-life is held constant at: A, B diffusion-specific estimates for mRNA export delay; C, D half of estimated mRNA export delays; E, F a quarter of estimated mRNA export delays.

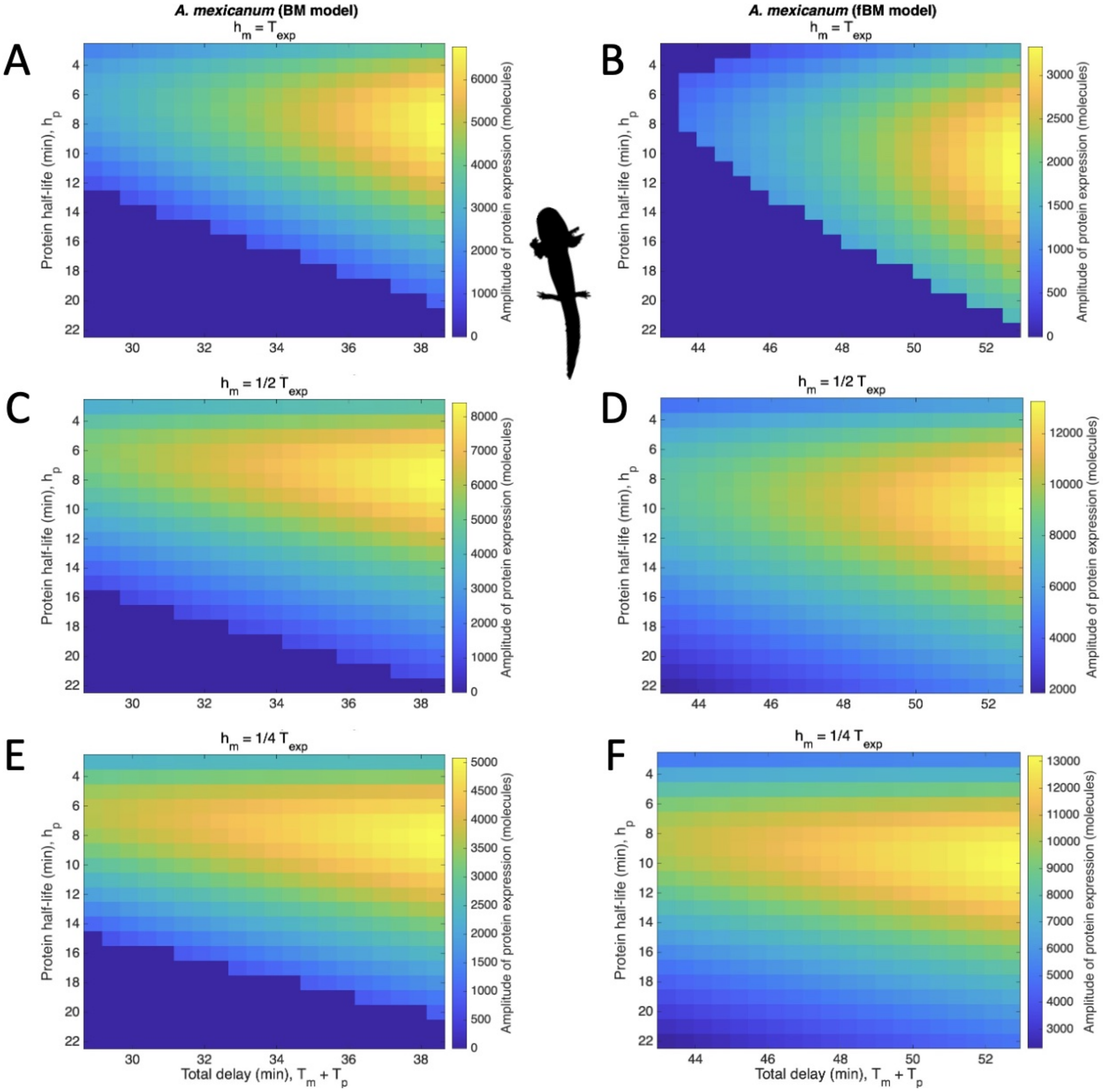


Figure S7: Resulting amplitudes of protein expression for *A. mexicanum* models corresponding to plots in figure 2. A, C, E normal diffusion/Brownian Motion model results; C, D, F obstructed diffusion/fractional Brownian Motion model results. mRNA half-life is held constant at: A, B diffusion-specific estimates for mRNA export delay; C, D half of estimated mRNA export delays; E, F a quarter of estimated mRNA export delays.
